# Molecular flexibility of high molecular weight hyaluronic acid measured by NMR has a profound effect on invasion of cancer cells

**DOI:** 10.1101/2024.12.17.626583

**Authors:** Uliana Bashtanova, Agne Kuraite, Rakesh Rajan, Melinda J Duer

**Author notes:** **Corresponding author:** Yusuf Hamied Department of Chemistry, University of Cambridge, Lensfield Road, Cambridge, CB2 1EW, UK. These authors are joint first authors (Uliana Bashtanova, Agne Kuraite).

## Abstract

Extracellular hyaluronic acid (HA) has been shown to be important in cancer; low molecular weight HA typically correlates with cancer progression, high molecular weight HA with homeostasis. Here we show that even high molecular weight HA can induce cancer cell migration when it is highly diluted. HA-induced cell signalling is primarily through HA binding to the cell surface receptor, CD44. We show by NMR spectroscopy that at high dilution, high molecular weight HA molecules access the conformations needed for strong binding to CD44 on the tens of nanosecond timescale, the relevant timescale for induction of CD44 signalling. We further show that, in contrast, at higher concentrations, HA molecules have insufficient flexibility for strong CD44 binding. The high dilution HA condition correlates with profound changes in brain cancer cell morphology and proteome which supports cancer cell invasion. We hypothesize that the flexibility of HA molecules is central to HA-mediated cell signalling and that this concept can explain previous observations that the outcome of HA-mediated signalling depends on the HA molecular weight. HA dilution leading to stronger HA signalling may be important in understanding the role that oedema plays in cancer recurrence after primary surgery.

## Introduction

Hyaluronic acid (HA) is a ubiquitous component of the extracellular matrix (ECM) in all tissues. Its deceptively simple polysaccharide structure belies its dual importance in the mechanical properties of the extracellular matrix^1^ and in determining cell fate.^1–7^ ECM HA is increasingly implicated in cancer progression.^8–12^ A range of studies have collectively led to the view that, in broad terms, high-molecular weight HA (HMW-HA; molecular mass > 500 kDa) typically supports normal tissue homeostasis and inhibits tumour formation whilst low molecular weight HA (LMW-HA; molecular mass 10 – 250 kDa) and HA oligomers (< 10 kDa) plays significant roles in cancer progression; however, there are numerous exceptions to these trends.^8,12–15^

HA activates cell signalling by binding to its primary cell surface receptor, CD44^16,17^ and also to other cell surface receptors including RHAMM, LYVE1, HARE/ stabilin2 and Toll-like receptors. Structural studies have shown that HA - CD44 binding requires both HA and the CD44 HA binding domain (HABD) to adopt specific conformations^18–20^ (Fig 1). The conformation of the portion of HA bound to the CD44 HABD in the postulated strong-binding mode^18^ (Figure 1C) is significantly distorted from the ribbon structures predicted to exist in bulk HA gels and solutions ^21–25^ (Figure 1B). Molecular dynamics studies suggest that the process of adapting the combined molecular conformations for binding takes of order hundreds of nanoseconds.^26^ Taken together, these features of the HA-CD44 HABD interaction suggest that the degree and timescale of conformational flexibility of HA may be significant in determining the rate of successful binding events and thus in initiating cell signalling, regardless of the HA molecular weight.

**Figure 1:**
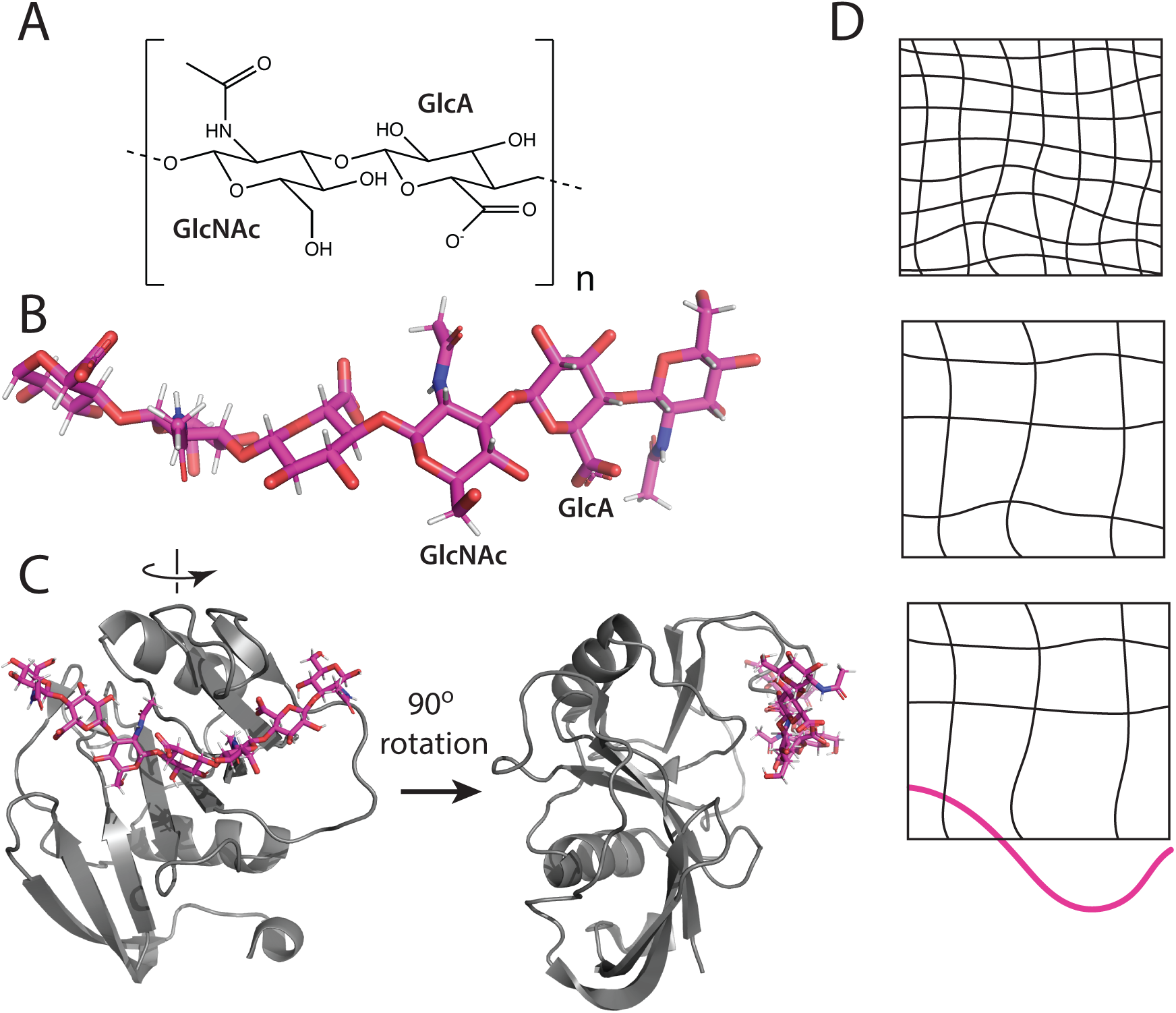
**A, B.** Molecular structure of HA is a polymer of repeating disaccharide N-acetyl-D-glucosamine (GlcNAc) and D-glucuronic acid (GlcA) rings. **C.** Type A binding of HA 8-mers on the CD44 hyaluronic acid binding domain (HABD) (from PDB 2JCQ^18^). The HA molecule must contort its conformation in order to access the HABD binding site in this ligand binding mode, and similarly in type B ligand binding (not shown, PDB 2JCR^18^). **D.** Schematic illustrating how reducing the HA concentration from 5 wt% to 2 wt% decreases the density of intermolecular contacts and entanglements and increases the probability of flexible HA segments able to bind to the CD44 HABD.

The conformational flexibility of an HA molecule in a pure HA gel is governed by the extent of polymer entanglement and the density and strength of its interactions with other HA molecules (Figure 1C,D). For a given HA concentration, shorter (lower molecular weight) HA molecules have lower degrees of entanglement than longer HA molecules (higher molecular weight) and so have greater flexibility. The same relationship between molecular weight and HA molecular flexibility will hold in an HA-rich ECM *in vivo* and, we reasoned, may be what underlies the observations of LMW-HA leading to cancer progression and HMW-HA tending to promote homeostasis; HA molecular weight may actually be a surrogate for molecular flexibility. If molecular flexibility is the fundamental parameter governing HA-CD44 binding and signalling, then HA concentration must also be an important factor. Dilution of an HA-rich ECM, for instance through oedema, will reduce both the density of intermolecular contacts and the degree of polymer entanglement, and increase HA flexibility, both in terms of the conformational degrees of freedom of the HA molecules and the timescale of their conformational interconversions. Thus, for a given HA molecular weight, the extent of hydration of an HA-rich ECM may be a determining factor in the probability of HA-mediating signalling from the ECM, suggesting that at sufficiently high dilution (increased odema), HMW-HA could also promote cancer progression instead of homeostasis.

Native brain ECM is an HA-rich ECM; it is abundant in HMW-HA and proteoglycans whose molecular structure mimics that of HA^27^ whilst fibrous ECM proteins such as fibronectin and collagens have only very low concentrations and are primarily confined to vessel walls, e.g. ∼ 1 wt% collagen IV in fresh brain.^27^ In a brain tumour setting such as glioblastoma multiform (GBM), oedema is common and so dilution of the brain HA can be expected to be relevant in GBM. We hypothesized that if HA flexibility is the important parameter in HA signalling rather than HA molecular weight, then sufficient dilution of the normal HMW-HA in brain ECM (or tumour-associated HMW-HA) would create HMW-HA molecules sufficiently flexible to induce CD44 signalling and GBM tumour progression, whilst brain ECM at normal dilution drives tumour homeostasis.

Thus, here we ask the question: Can excessive hydration of an HA-rich ECM lead to increased HMW-HA polymer flexibility and drive altered brain cancer cell signalling?

We chose a simple 3D model of brain ECM of varying concentrations of HMW-HA solubilised in cell culture media so that we could be sure any differences in cell behaviour were a result of HA concentration. Normal brain ECM has ∼ 3 wt% HA plus ∼ 4.5 wt% GAGs,^27^ which mimic HA in chemical properties. To model the polyanionic hydrogel environment of normal brain, we thus used HMW-HA to account for both HA and GAGs in the brain ECM and chose HMW-HA concentrations which resulted in HA-cell culture media gels with rheological properties (storage and loss modulus) similar to healthy human brain^28,29^ (5 wt% HMW-HA), with a storage modulus of order one order of magnitude lower (2 wt%), and one order of magnitude higher (10 wt%) than normal brain (see Fig S1 for rheology data).

The molecular conformational changes important for CD44 binding and signalling occur on the 10s – 100s of nanoseconds timescale according to molecular dynamics studies.^26,30,31^ Nuclear magnetic resonance (NMR) ^1^H spin-lattice relaxation time constants, T_1_, give highly useful information on nanosecond timescale molecular motions. The ^1^H T_1_ for water-binding polymers like HA depend three terms: (i) molecular conformation changes or rotations on the nanosecond timescale; (ii) cross-relaxation between the dipole-coupled ^1^H in the polymer through spin diffusion, and (iii) ^1^H exchange between exchangeable ^1^H in the polymer and surrounding water molecules^32–35^ (see SI for equations).

In rigid and semi-rigid polymer networks where individual molecular motion with nanosecond correlation times is strongly restricted, term (i) is small, and the cross-relaxation of term (ii) is expected to dominate (see SI, equation (1)). This scenario is identified by all the observed polymer ^1^H signals having the same T_1_ relaxation time constants. In contrast, where in a situation where individual polymer molecules have molecular conformational flexibility with nanosecond correlation times, term (i) makes a significant contribution and term (ii) is quenched (because the ^1^H-^1^H dipolar coupling that drives this term is partially averaged by the molecular motion). In this scenario, the ^1^H T_1_ time constants depend on the degree of nanosecond timescale motion of the different polymer functional groups containing the ^1^H; typically, methyl groups can rotate rapidly compared to bulkier groups, like the sugar rings that comprise the HA backbone. Thus, molecular flexibility on the nanosecond timescale is indicated by methyl and sugar ring ^1^H having different T_1_, methyl ^1^H typically having smaller T_1_ relaxation time constants.

Term (iii) couples the relaxation of the polymer ^1^H and that of the water ^1^H, which results in the polymer (and water) ^1^H relaxation being a sum of two exponential processes with time constants T_1_^+^ and T_1_^−^ that are weighted averages of the true polymer and water ^1^H T_1_ from which the HA and water ^1^H T_1_ can be extracted (see SI, equation (6)).

Figure 2A shows the ^1^H T_1_ measured at 37°C for HMW-HA methyl groups in the N-acetyl-D-glucosamine rings and the non-exchangeable sugar ring ^1^H collectively for HA dilutions of 10, 5 and 2 wt% in cell culture medium. The ^1^H T_1_ are the same within error for methyl and ring ^1^H at the higher HA concentrations (5, 10 wt%), indicating that ^1^H cross relaxation, spin diffusion, is the dominant term governing HA ^1^H spin lattice relaxation at these higher concentrations. In contrast, the methyl and ring ^1^H T_1_ are significantly different to each other in 2 wt% HMW-HA, showing that spin diffusion is no longer the dominant term governing relaxation and that thermal molecular motions now govern the ^1^H T_1_. These data are consistent with rheology measurements on the HA gels (Fig S1) which show that the crossover oscillation frequency for the storage and loss moduli which represents the intrinsic molecular disentanglement rate is an order of magnitude higher for the 2 wt% HA gel compared to 5 wt%.

**Figure 2:**
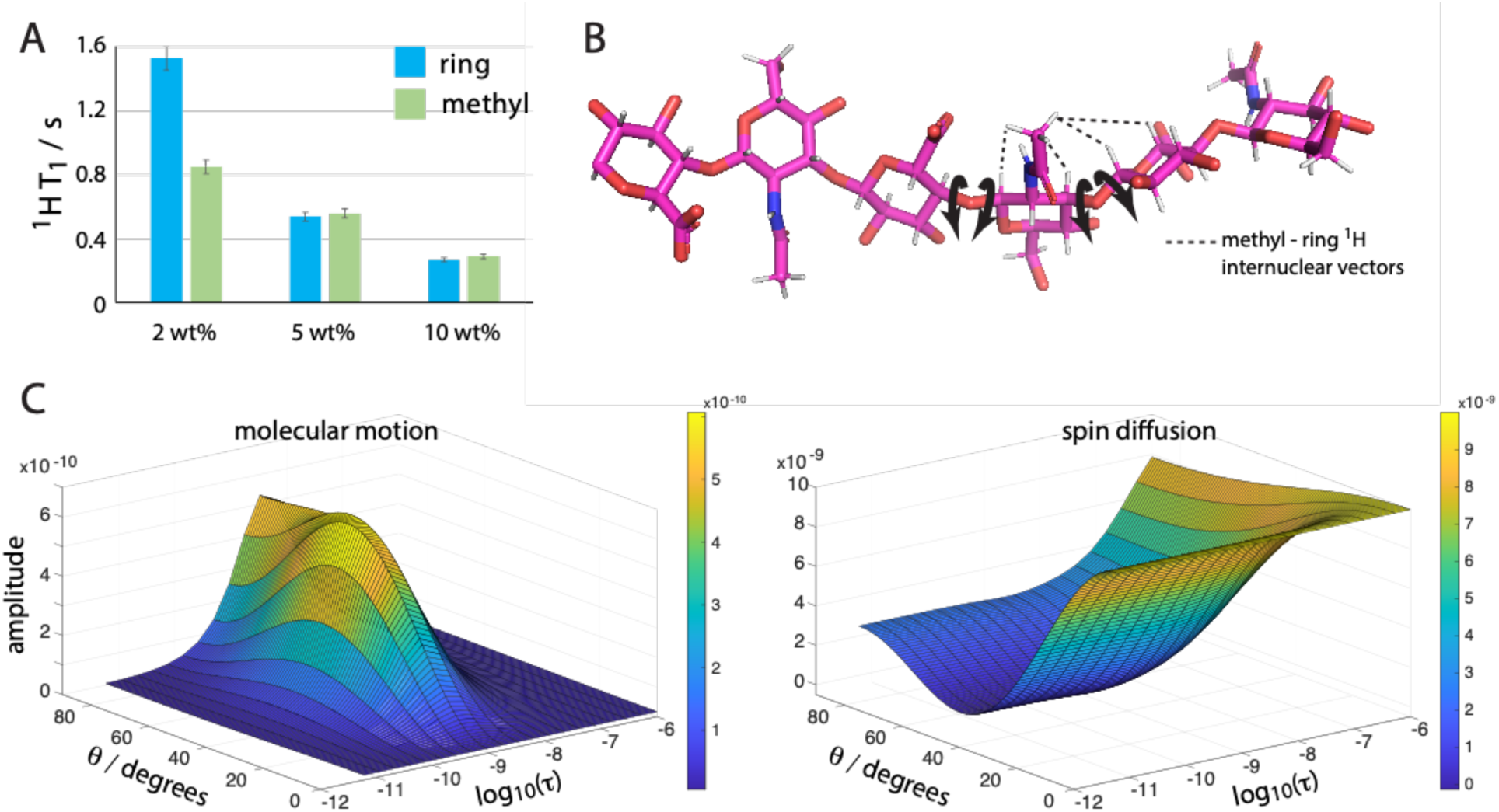
**A.** ^1^H T_1_ data for HMW-HA ring (non-OH) and GlcNAc methyl ^1^H as a function of HA concentration. See SI for details of T_1_ data analysis and Fig S2. **B.** Schematic illustrating the HA internal molecular motions around the glycosidic bonds that contribute to ^1^H T_1_ relaxation. Motions that change the orientation of ^1^H-^1^H vectors can potentially contribute to ^1^H T_1_ for HA. **C.** The amplitudes of the contributions of internal HA molecular motions with correlation time τ and spin diffusion in the presence of isotropic molecular tumbling (with correlation time τ_macro_ = 10^−7^s here; see Fig S3 for simulations with other τ_macro_) to ^1^H T_1_ See SI for simulation details. The internal motion is random ^1^H-^1^H vector motion on a cone of cone angle 8. The simulation shows that the molecular motion term dominates over the spin diffusion term only for τ between ∼10^−8^ s and 10^−10^ s and angular amplitudes of motion above ∼30°.

To understand the range of thermal motion correlation times and angular amplitudes of that motion that result in the ^1^H spin-lattice relaxation molecular motion term dominating over the spin diffusion term, we modelled how the size of these two terms vary with correlation time and motional amplitude. The dominant interaction leading to ^1^H relaxation in the HA system is ^1^H-^1^H dipolar coupling. Fluctuations with time of this dipolar interaction through molecular motions that alter the orientation of ^1^H-^1^H vectors drive the ^1^H spin-lattice relaxation. We modelled the case for random motion of a ^1^H-^1^H vector on a cone of cone angle θ (amplitude of the molecular motion). The molecular motion of the HA rings and side groups is clearly highly complex and will consist of a spectrum of motional modes about different bonds and molecular axes with different amplitudes and correlation times. Our aim is to understand the order of magnitude of the angular amplitudes of motion and correlation times that would lead to the molecular motion term governing HA ^1^H T_1_ to dominate over the spin diffusion term. Hence, we chose this simple model of molecular motion, knowing that the true molecular motions can be considered as made up of contributions from similar simple motions about (many) individual bonds and molecular axes. The results from our model in Figure 2C show that the spin diffusion term has a maximum value that is approximately an order larger than the molecular motion term and thus will always dominate unless the molecular motion meets the conditions for being close to maximising the molecular motion term; that is (from Figure 2C) correlation times between 10^−8^ s and 10^−10^ s and angular amplitudes of motion above ∼30°.

Previous molecular dynamics calculations have indicated that HA oligomers need to adapt their conformation to the CD44 HABD binding site geometry on the tens to hundreds of nanoseconds timescale,^26^ whilst previous analysis of crystal structures for free and CD44 HABD-bound HA shows a kink of around 22° in the HA molecular long axis is needed for HABD binding (see Fig S4).^18^ Our ^1^H NMR T_1_ analysis shows that HMW-HA polymers have this timescale of molecular motion when in low concentration (2 wt% here) but not at higher HMW-HA concentrations (here 5 wt% and higher) and moreover, at this low HMW-HA concentration, the HA polymers have thermal amplitudes of angular motion of at least the 22° needed for HA to bind to the CD44 HABD. We hypothesized therefore that 2 wt% HMW-HA could have a different signalling effect on cancer cells than 5 wt% HMW-HA where HA polymers have relatively little flexibility on the nanosecond timescale.

Thus, we next modelled *in vitro* how brain cancer progresses and invades as a function of HMW-HA concentration in brain matrix. We generated 3D spheroids from two cancer cell lines of neuroepithelial origin (U251 and U87) and also SP20 tumour-initiating cells isolated from patients glioblastoma tumour tissue after surgical resection.^36^ We placed these spheroids in a range of HMW-HA concentrations to exemplify the cases of flexible and stable HA networks as discovered from our NMR analysis and which have comparable stiffness to the range of brain ECM stiffness in cancers (see Figure 2D).^37^ We found that for all cell types, invasion into 2 wt% HMW-HA was very advanced by day 4 (Figure 3), but only a few cells were dispersed into 5 wt% HMW-HA (Figure 3) and their quantity did not change for periods of up to 6 weeks. Cell morphology was also different: in 5 wt% HMW-HA cells were typically rounded and lacked invadopodia, but in 2 wt% HMW-HA cells adopted spindle or star-shape flattened morphologies with prominent invadopodia (Figure 4A). In a control liquid medium, cancer spheroids demonstrated variable behaviour from a full dispersion of U251 cells to almost no dispersion of U87 cells (Figure 3). Thus, at least for the U87 cell line it is the concentration/flexibility of HMW-HA which controls invasion, rather than the viscosity of the environment. To gain some understanding of how HMW-HA flexibility controls U87 cell behaviour, we utilised proteomic approach as we reasoned that the radical difference in cell morphology and behaviour in 2 versus 5 wt% HMW-HA had to be underpinned by widespread changes in protein biosynthesis.

**Figure 3.**
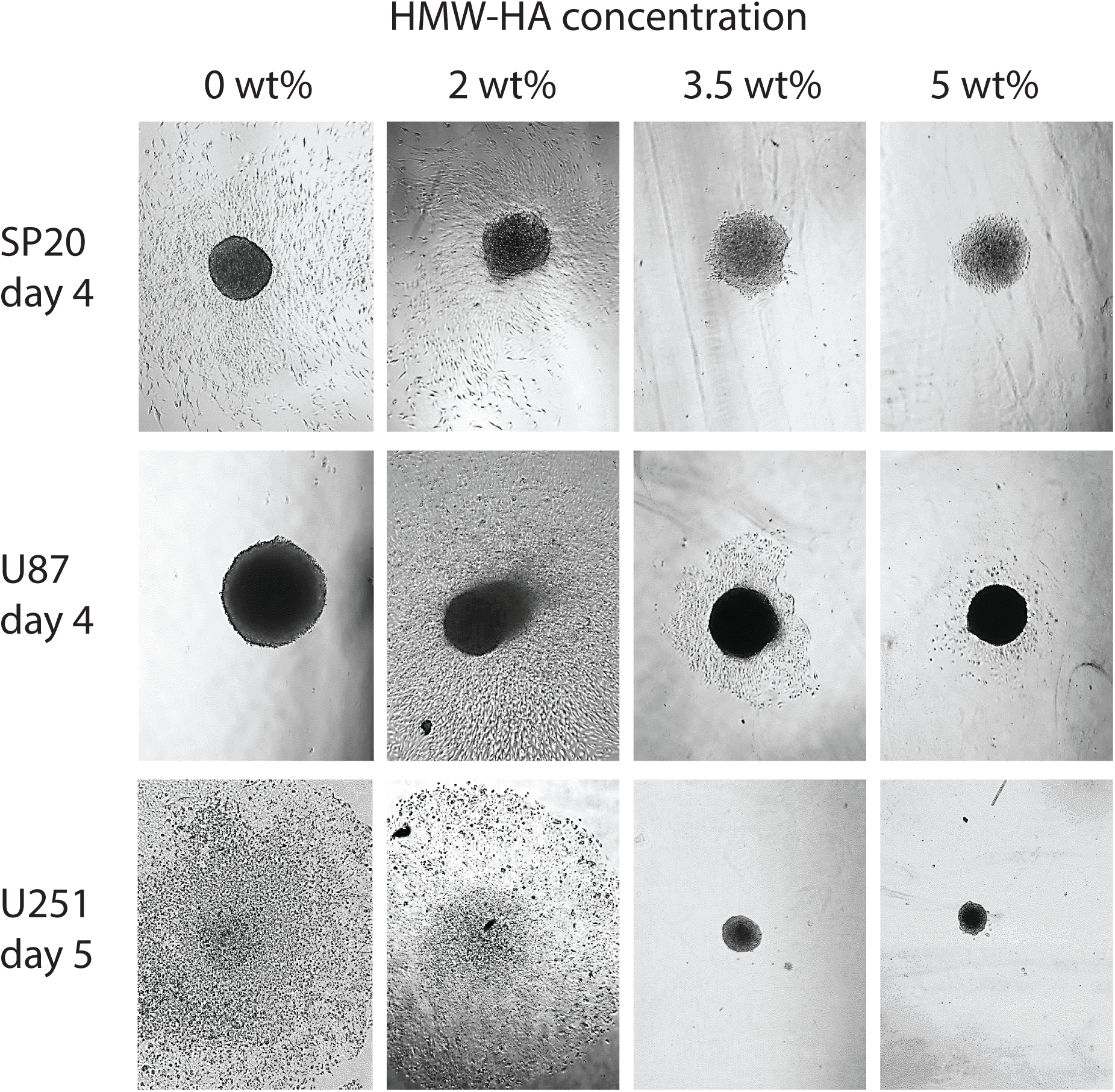
Cancer spheroid behaviour in different HMW-HA concentrations. Spheroids were grown in media for a period of 7 days, after which multiple spheroids were embedded in each concentration of HMW-HA media gel. Representative microscope images of cell migration from the spheroids after 4 or 5 days of culture in different HA concentrations illustrate typical spheroid behaviour in each specific HMW-HA concentration. Field view 1900µm x 2500µm.

**Figure 4.**
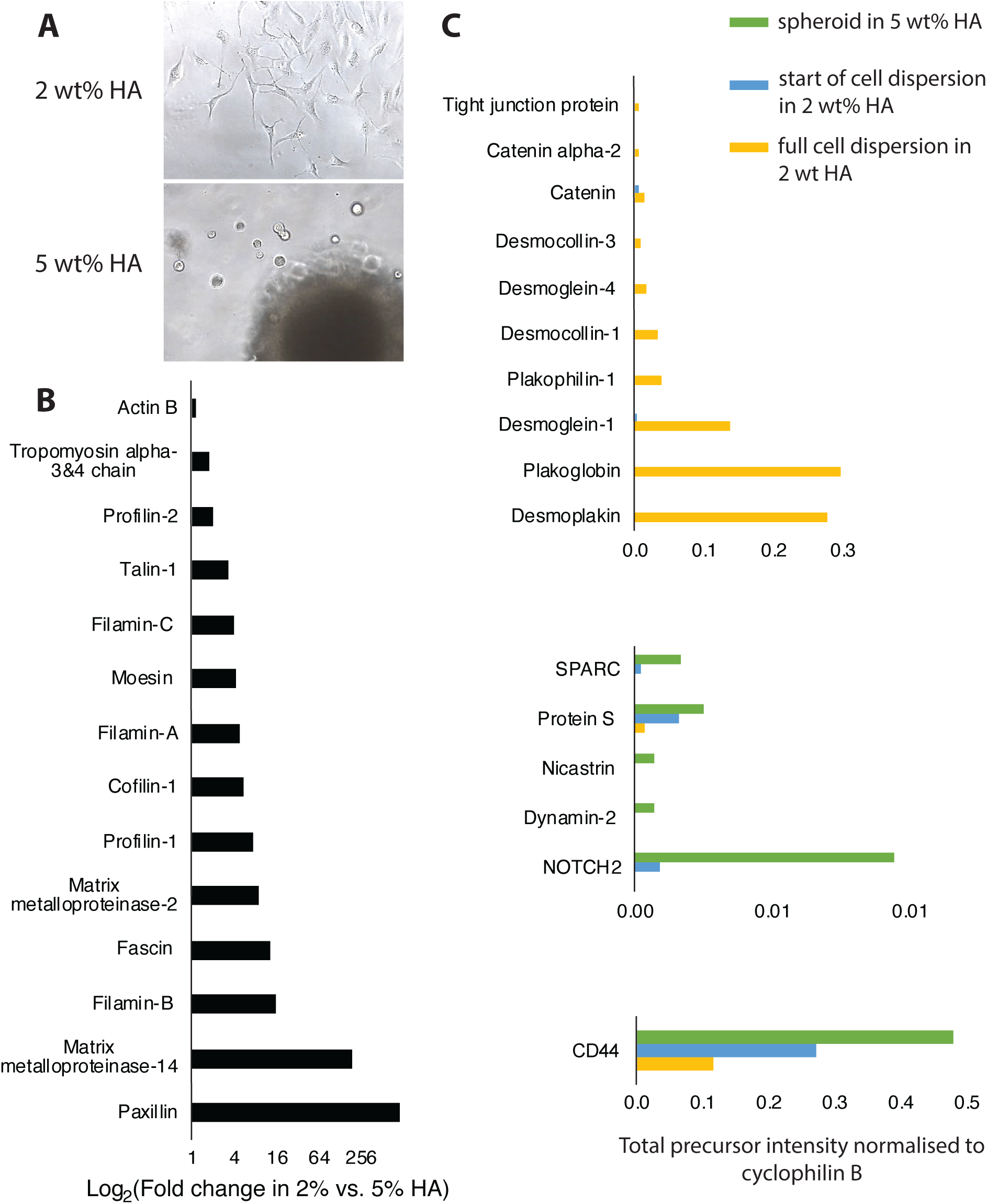
Changes in U87 proteome relevant to changes in cell morphology and fate in HMW-HA. **A.** Morphology of U87 cells dispersed from spheroid in HMW-HA. Field view 1900µm x 2500µm. **B.** Upregulation of invadopodium proteome in 2 wt% HMW-HA after full cell dispersion from spheroid compared to spheroid in 5 wt% (sample 2 and 1, respectively, see Experimental Methods). **C.** Changes in cell-cell junction proteome and CD44/Notch axis in different HMW-HA concentrations and at two different stages of cell dispersion in 2 wt% HMW-HA. Green – spheroid in 5% HMW-HA (sample 1), blue - beginning of cell dispersion in 2% HA (sample 3), yellow – full cell dispersion in 2% HA (sample 2)

Shotgun label-free proteomics is widely used in discovery studies. For our pilot study, we used it in a semi-quantitative mode (i.e. without extensive replicas), so it was important to minimize any contributions to variability unrelated to change in HMW-HA concentration. The contribution to variability in human brain samples was previously assessed by Piehowsky et al. ^38^ and rated as follows: sampling of biological tissue (72%) ≫ instrumental variance, i.e. short-term run-to-run instrumental response fluctuation (16%) > instrumental stability, i.e. long-term drift of the quantitative response of the LC–MS/MS platform over a 2 week period (8.4%) > digestion (3.1%)^38^. However, for *in vitro* cell cultures it was shown that total coefficient of variation (CV) was much lower compared to tissues: CV for bacterial culture was 11%^39^ compared to CV = 34% for *ex vivo* human brain samples.^38^ Thus, using *in vitro* grown U87 cancer cells permitted implementation of semi-quantitative approach to compare protein expression in samples grown at different HA concentrations with the caveat that all precautions were taken to minimise variability arising from instrumentation, sampling and protein extraction and digestion (see Experimental Methods). Furthermore, in our analysis we decided to rely on directional changes in groups of structurally-co-operating proteins, for example, in the invadopodium scaffold proteome and cell-cell junction proteome.

To corroborate this approach, we first analysed the group of invadopodium scaffolding proteins, because this cellular structure was clearly present in 2 wt% HMW-HA but absent in 5 wt% HMW-HA. We found that key invadopodium scaffold proteins were all upregulated by 2 to 200-fold in 2 wt% compared to 5 wt% HMA-HA (Figure 4B). Foremost upregulated were proteins that structurally reorganise actin into stable filaments and bundles (Figure 4B) such as fascin-1, which ensures actin bundle stability by binding to F-actin at regular intervals,^40^ filamins A and C, known to be involved in non-covalent cross-linking of F-actin,^41^ and tropomyosin 3 and 4, as a master regulator of individual F-actin filament function, with isoforms 3 and 4 most characteristic for invasive cancers.^42^ Proteins implicated in early stages of invadopodium formation were also upregulated (Figure 4B), including profilins which promote actin assembly when barbed ends are free^43^ and cofilin-1 which drives membrane protrusion by severing actin filaments and their directional extension by treadmilling.^44^ Also upregulated were three key proteins, which connect the internal actin cytoskeleton to adhesion receptors on plasma membrane (Figure 4B), in particular moesin which links actin filaments to CD44,^45^ and talin and paxillin which are essential for activation of integrins,^46^ recruitment of phosphatases and kinases and their interconnection with actin bundles.^47^ The invadopodium is considered mature when it recruits matrix metalloproteases (MMPs),^48^ and in 2 wt% HMW-HA, MMP2 and MMP14 were highly upregulated compared to 5 wt% HMW-HA (Figure 4B). Expression of β-actin did not change in different HMW-HA concentrations, as expected (Figure 4B), because β-actin is normally reorganised in different environments rather than regulated by biosynthesis^49^ and hence widely used as an internal standard and ‘housekeeping control’ in protein studies.

Another noticeable cell morphological feature, which was present in 2 - 3.5 wt% HMW-HA, but absent in 5% HMW-HA, was a collectively moving ‘cellular sheet’ (Figure 5B). It was previously shown that collective cell motility, i.e. mass migration of cancer cells in sheets, strands and clusters, results in a very rapid invasion in comparison with individual cell movement, but requires maintenance of cell-cell junctions.^50^ For example, elevated expression of desmoplakin and plakoblobin – the two most abundant proteins in desmosomal junction - facilitate breast and lung cancer cells to form clusters and increase their survival in circulation.^51^ In 5 wt% HMW-HA, no proteins from any type of cell-cell junction were detected (Figure 4C), but in 2 wt% HMW-HA, all desmosomal cadherins and linker proteins were present with desmoplakin and plakoglobin being the most upregulated (Figure 4C). Proteins from other junctions were also present, for example α-catenin from adherens junction and ZO-1 from tight junctions, but only the desmosomal proteome was complete (Figure 4C). Hence, we can speculate that the ‘cellular sheet’ observed in 2 - 3.5 wt% HMW-HA (Figure 5B) was held together primarily by desmosomes.

**Figure 5.**
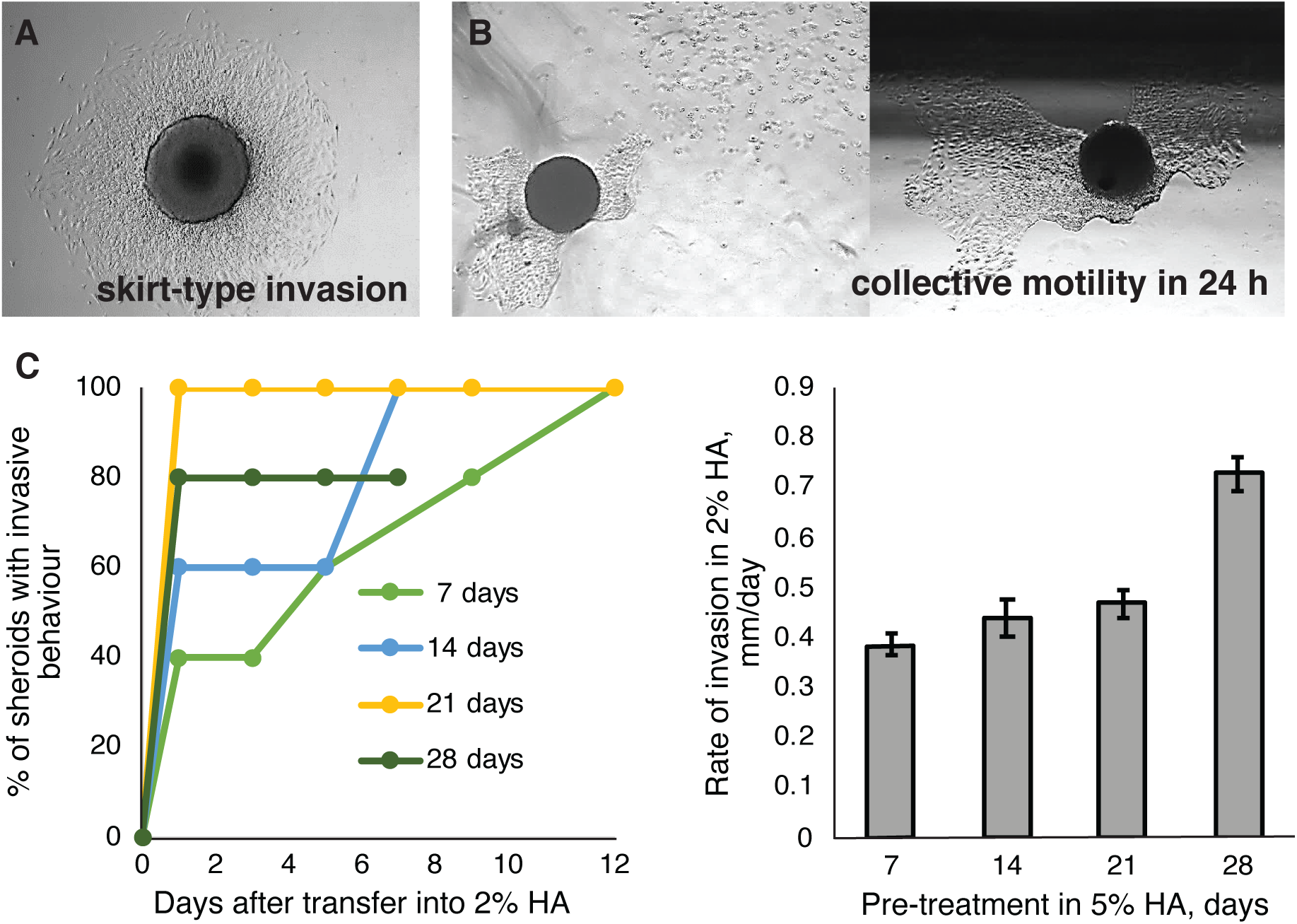
U87 spheroid dormancy and invasiveness in HMW-HA. Spheroids were cultivated in liquid medium for 7 days and placed into HMW-HA. **A.** Typical skirt-type invasion inside 2 wt% HMW-HA. Invasion occurred on day 1 after spheroid placement and continued as a gradual increase in ‘skirt’ radius. By day 7-10 spherical structure became invisible, i.e. full cell dispersion was achieved. Field view 1900µm x 2500µm. **B.** Collective cell motility inside 2-3.5 wt% HMW-HA. Sometimes a cellular sheet of an irregular form appeared around spheroid on the 2-3 day after spheroid was placed in HMW-HA; after 24 h spheroid and cellular sheet were found several mm away from their initial position, for example, next to the well wall (black shadow on the top of right picture). Dispersion of single cells was also observed, and their movement was independent from the cellular sheet and spheroid. Field view 1900µm x 2500µm. **C.** Invasion into 2 wt% HMW-HA after a period of dormancy in 5 wt% HMW-HA. Spheroids were placed inside 5 wt% HMW-HA for 7, 14, 21 and 28 days (five spheroids per each period of dormancy). During dormancy period spheroid shape and diameter did not change, and no cell dispersion was observed. When spheroids were transferred into 2 wt% HMW-HA percent of spheroids able to disperse cells (‘invasive spheroids’) was calculated (left). Rate of invasion was calculated as an enlargement ‘skirt’ radius per day (right). Error bars represent SE.

Based on the expected changes observed in invadopodium and cell-cell-junction proteome at different HMW-HA concentrations, we considered these samples representative for a pilot detection of proteins which underpin physiological changes in U87 cells. The major physiological change which occurred in 5 wt% HMW-HA compared to 2 wt% HMW-HA was spheroid dormancy: cancer spheroids placed in 5 wt% HMW-HA did not grow, invade or die, but remained dormant until transferred into to 2 wt% HMW-HA (Figure 5C). In 5 wt% HMW-HA, the expression of 193 proteins was increased > 4-fold and 138 proteins were synthetized *de novo* compared to 2 wt%-HMW HA (see Table S2), thus are likely to be involved in quiescence uphold in U87 cells. According to GO annotation, these proteins are involved in biological processes essential for cell survival and metabolism support, such as: chromosome packaging, ribosome hibernation and biogenesis, mitochondria maintenance, cellular response to starvation, apoptosis suppression, tumor acidosis, glucose storage, activation of hexosamine pathway, peptide metabolism, protein biosynthesis and control of folding, proteolysis regulation, and protein translocation and trafficking (see Table S2).

Substrate mechanics affects cell fate transition from proliferation to dormancy,^52^ and is coordinated by intracellular signalling with the Notch pathway being widely recognised as a major determinant of cell fate across all metazoans.^53^ In our data, Notch-2 was strongly upregulated for spheroids in 5 wt% HMW-HA and below detection in 2 wt% HMW-HA after total cell dispersion from the spheroid (Figure 4C). Notch-1 was not detected in either of the samples. Hence, similar to adult brain where Notch-2 was shown to control quiescence of neuronal stem cells,^54,55^ Notch-2 may convey quiescence of U87 cells in 5 wt% HMW-HA. Search for an HA receptor in our model yielded only one protein – CD44, which was upregulated following the increase in HMW-HA concentration as expected for a chemoreceptor (Figure 4C). The CD44 and Notch signalling pathways share two key proteins: γ-secretase, which cleaves intracellular domains known as CD44-ICD and NICD respectively^56–58^ and dynamin, which enables formation of endocytosis vesicles essential for nucleus translocation of both domains.^59,60^ Indicative of activation of the CD44/Notch axis during dormancy in 5 wt% HMW-HA, dynamin-2 and nicastrin (part of the γ-secretase complex) were detected for spheroids in 5 wt% HMW-HA, but not in 2 wt% HMW-HA (Figure 4C). Two regulators of the Notch signalling pathway – osteonectin (SPARC) and vitamin K-dependent protein S (PROS1)^61,62^ were also upregulated in 5 wt% HMW-HA (Figure 5C), again indicative of Notch involvement in U87 cell fate determination in 5 wt% HMW-HA.

Given that the only difference between spheroids in 2 and 5 wt% HMW-HA initially is the HA concentration and that the only abundant cell surface receptor for HA in our proteomics data was CD44 (Figure 5C), the differences in cell morphology and behaviour (Figure 3, 4, 5) are likely to be the result of differences in HA-CD44-mediated signalling and correlate with the nanosecond timescale conformational flexibility of the HA polymers discovered by NMR spectroscopy (2 wt% HMW-HA) or relative lack of such molecular flexibility (5 wt% HMW-HA). This supports our hypothesis that CD44 signalling is induced by HA molecules that are flexible on the nanosecond timescale and that in turn depends on whether the HA molecules are strongly or weakly bound into the ECM.

We reasoned that if HA molecules strongly bound into the ECM inhibit CD44 signalling then chemically-crosslinking HA into the ECM, to substantially increase the binding strength of HA into the ECM, should result in similar inhibition of CD44 signalling. To test this, we synthesized a crosslinkable HA polymer, oxidized hyaluronic acid (oxHA) where 35% of the glucuronic resulting in two aldehyde groups on the oxidized ring (see SI for details of oxHA characterization). Hence, most of the oxidized hyaluronic acid molecule has the same composition and structure as native hyaluronic acid, but its aldehyde groups on the oxidized glucuronic acid rings will rapidly crosslink to terminal amine groups of proteins in the ECM and thus significantly increase its strength of binding to the ECM.

We then repeated our cell migration study and proteomics measurements for U87 spheroids in a 50: 50 by weight mixture of 2 wt% HMW-HA and 2 wt% oxHA. The oxHA has a lower molecular mass than the original HMW-HA (see SI) which would be expected to drive cancer cell growth and/ or migration if HA molecular mass were the only HA parameter important in cell signalling. However, we found that there was no observable U87 cell migration or spheroid enlargement in the oxHA-HA mixture, in contrast to the case of 2 wt% HA alone, where cell migration was rapid and substantial (Figure 3).

We hypothesised that the arrest of invasion caused by oxHA was the loss of conformational flexibility in HA-chains due to cross-linking to ECM proteins, which in turn would cause changes in protein expression similar to that for cells in 5 wt% HMW-HA rather than in 2 wt% HMW-HA. Indeed, we found that ox-HA treated cells expressed Notch-2 and Notch signalling pathway proteins as in 5 wt% HMW-HA (Table 1). Furthermore, ADAM proteases involved in extracellular cleavage of Notch and sugar transferases required for Notch glycosylation^63^ were detected in oxHA sample, implying that regulation of cell fate in oxHA may depend on the Notch pathway even more.

**Table 1.**
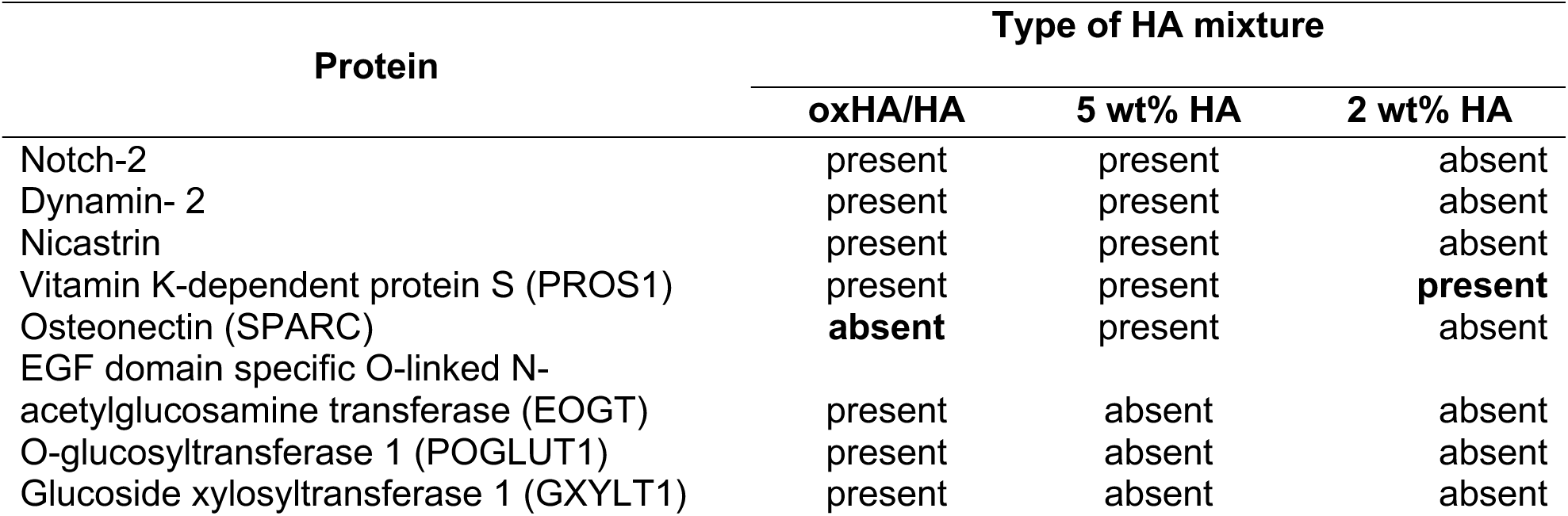
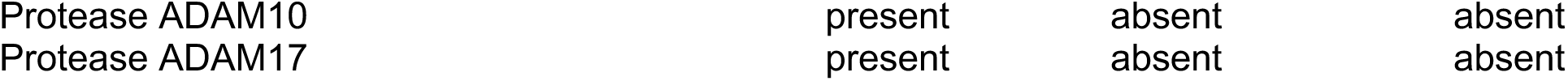
Detection of proteins involved in Notch signally pathway in 2 wt% HMW-HA, 5 wt% HMW-HA and oxHA/HA mixture by shotgun label-free proteomics.

## Discussion

### HA flexibility and dissociability on the nanosecond timescale is required for HA-CD44 induced signalling

HA-CD44 signalling is initiated by close intermolecular contact (binding) of an HA molecule segment ∼6 - 8 saccharide rings in length within the binding pocket in the CD44 HABD.^18,19,26,30^ For this binding to occur, several processes have to happen: (1) The relevant segment of the HA molecule must detach from its ECM intermolecular contacts (Fig 1D). (2) The binding segment must distort from its typical ECM conformation.^18^ These two processes must occur on a time scale which is faster than the typical time taken for the CD44 HABD to change its conformation to allow strong HA binding, which from molecular dynamics calculations is a few 100 nanoseconds.^26^ To initiate invasion signalling, the resulting CD44-HA complex must interact with the relevant members of the invadopodia development pathway. For these interactions to occur, CD44 must be released from cholesterol-rich lipid raft in the plasma membrane into phosphatidylinositol 4,5-bisphosphate (PIP2)-enriched region.^64^ PIP2 activates the phosphorylation of CD44 and, in turn, that phosphorylation allows CD44 to bind to ERM proteins (moesin/ezrin/radixin) which activates the CD44-dependent Rho-signalling pathway and invadopodia extension.^65^ Binding a hydrophilic HA molecule to CD44 residing in hydrophobic lipid raft can be expected to lower the energy for translocation into PIP2-enriched (hydrophilic) membrane regions, but translocation also requires either that the HA molecule is sufficiently flexibly to allow the CD44 HABD – HA binding to persist as CD44 to moves between membrane domains, or the CD44-bound HA must readily dissociate from its ECM contacts to allow the translocation. Thus, HA flexibility and its strength of binding into the ECM are highly relevant both for the initial HA binding to the CD44 HABD and for the subsequent signalling pathways.

### NMR shows that low concentrations of HA allow HA to access the required geometries for binding on the appropriate timescale

We have shown here using NMR ^1^H T_1_ relaxation times (Figure 2) that a low HMW-HA (∼ 1MDa) concentration (2 wt%) achieves the HA thermal flexibility requirements for CD44 HABD binding^18^ namely HA ring rotational motional modes with amplitudes of at least 22° with correlation times of 10s of nanoseconds.^66^ That same low HMW-HA concentration (2 wt%) induces aggressive cell invasion for three different brain cancer cells lines (Figure 3), consistent with other recent work which showed that a wide range of patient-derived GBM cells *in vitro* can be induced to invade their surrounding ECM if the HMW-HA concentration is sufficiently low.^67^ In contrast, cancer cell spheroids in a HMW-HA concentration representative of the polyanion (HA + glycosaminoglycans) concentration in normal brain (5 wt%),^27^ show very little cancer cell migration from the spheroids or spheroid growth. Our NMR data show that in 5 wt% HMW-HA, there can only be much lower amplitude HA internal motion on the nanosecond timescale, well below the 22° needed for HA to bind to CD44 HABD (Fig 2C). The lack of invasion for one of the cell lines (U87) in the presence of only media, i.e. low viscosity medium, yet aggressive invasion in 2 wt% HMW-HA (Figure 3) suggests that cancer cell invasion with reduced HA concentration is not simply a viscosity effect. Our pilot proteomics data on the samples obtained from cells in 5% and 2 wt% HMW-HA clearly indicates that U87 cell proteome undergoes considerable regulation of protein expression in response to the change of molecular flexibility of HA between these two HMW-HA concentrations: cell morphology, cytoskeleton structure, cell-cell junctions and overall cell physiology. Our most important finding is a possible involvement of Notch-2 and its signalling pathway (dynamin-2, nicastrin, SPARC and PROS1) in dormancy of cancer spheroids in a HMW-HA concentration (5 wt%) representative of normal brain ECM. Hence, similar to adult brain where Notch-2 was shown to control quiescence of neuronal stem cells,^54,55^ Notch-2 may convey quiescence to brain cancer cells when HA molecular flexibility is low.

Increased HA molecular flexibility can be expected wherever there is a reduction in HA concentration through increased water content of a tissue or where the HA network is disrupted by other ECM components. Oedema as a result of brain tumour surgical resection is common and can be expected to dilute HA in the tumour margins potentially initiating invasion signals for remaining cancer cells; significantly, it is in these same regions where tumour re-growth is commonly seen. Infiltration of the brain extracellular HA/ glycosaminoglycan network by extracellular matrix proteins expressed by cancer cells or cancer-associated fibroblasts is another potential scenario that may increase the probability of HA dissociation from the ECM during their thermal motions, and so the conditions under which cell signalling is initiated. Interestingly, Safarians *et al*^67^ found that the HA concentration which which maximized patient-derived GBM cell migration was dependent on the cell line, whilst other work shows that HA interacts primarily with the cell glycocalyx and much more rarely with CD44.^68^ Taken together, these works suggest that HA – CD44 interactions are finely tuned by the cell surface and that the cell glycocalyx is likely to be an important further moderator of HA molecular flexibility.

HA degradation into lower molecular weight fractions is a commonly observed scenario in tumours^8,9,69–71^ and can also be expected to result in more flexible HA molecules because of the lower degree of polymer entanglement for shorter (lower molecular weight) molecules. HA degradation is frequently correlated with poor patient prognosis and disease progression,^69,70^ which we speculate is related fundamentally to the increased flexibility of lower molecular weight HA leading to a higher probability of strong HA – CD44 binding events, and concomitant CD44-mediated cell signalling, rather than being a direct function of molecular weight. Rather, HA molecular weight has become an inadvertent surrogate for molecular flexibility.

## Acknowledgements

Funded by the European Union (ERC, EXTREME 101019499). Views and opinions expressed are however those of the author(s) only and do not necessarily reflect those of the European Union or the European Research Council Executive Agency. Neither the European Union nor the granting authority can be held responsible for them. AK acknowledges funding from a UK EPSRC studentship, doctoral training allocation to the Yusuf Hamied Department of Chemistry, University of Cambridge. RR and UB were funded in part by a research grant from Cambridge Oncology Ltd to the University of Cambridge (1/1/2019 – 31/12/2022). UB was also funded by the ERC grant EXTREME 101019499.

The authors would like to thank Dr David G. Reid and Dr Thomas Kress for assistance with oxHA NMR, the Melville Laboratory for Polymer Synthesis, University of Cambridge for use of their rheology equipment and the Cambridge Centre for Proteomics, Department of Biochemistry, University of Cambridge for running LC-MS/MS and protein identification for this paper.

## Competing interests

Melinda J Duer is a shareholder of Cambridge Oncology Ltd.

## Experimental Methods

### Production of HA 3D matrix

A series of hyaluronic acid (HA) (Streptococcus equi, 91%. M_w_= 1MDa, Alfa Aesar, Thermo) 3D matrices with varying concentrations (2, 5, 10wt%) were produced either for measurements of cell/spheroid growth or NMR or rheology. For NMR the diluent was deuterium oxide (D_2_O) (99.8%, Sigma Aldrich), for rheology - MilliQ water (Millipore Advantage A10), for cells/spheroids - Minimum Essential Medium (MEM) ([-] Glutamine, [+] Earles salts) supplemented with L-glutamine–penicillin-streptomycin, Fetal Bovine Serum, and occasionally Amphotericin B. Hyaluronic acid was mixed with diluents and then kept in the incubator at 37°C, 5% CO_2_ and mixed periodically until it was fully dissolved, and then kept at room temperature for a few days before use. If needed, the viscose solutions of HA were centrifuged using Heraeus Instruments Biofuge 13 to remove bubbles at 9000 rpm, taking care to not precipitate the polymer, with 2wt % samples being spun for <30 sec or left to settle without centrifuging, 5wt % samples for 30 – 60 sec, 10wt % samples for 3 – 5 min.

### Synthesis of oxidized HA (oxHA)

The synthesis of oxidized hyaluronic acid (oxi HA) was based on that of Su et al. (2011),^72^ with specific modifications incorporated. Hyaluronic acid was dissolved in 100 ml of Milli-Q water to make a solution with a final concentration of 1 wt%, left overnight at room temperature. To oxidise the hyaluronic acid, sodium periodate (2.6%) in PBS was added dropwise to the solution while stirring. The reaction was allowed to proceed for 3 hours and 24 hours in the dark to achieve different levels of oxidation, after which it was stopped by adding 2.5 ml of 10% ethylene glycol. The solution was dialysed using 6-8 kDa dialysis tubing with Milli-Q water as the buffer. The dialysis lasted for seven days, with the water being replaced twice daily. After the dialysis process, the oxi HA solution underwent a volume increase of twice its original size due to water absorption. The solution was then transferred to a 50 ml tube and rapidly frozen using liquid nitrogen. Subsequently, it was subjected to freeze-drying in the Vir Tis freeze dryer for seven days. The freeze dryer was set to maintain a vacuum pressure of 100 mTorr and a condenser temperature of −76°C. See SI for product characterization.

### Cell culture

#### Materials

Accutase (Thermo), Alpha Eagle Medium (phenol red free) (Alpha MEM; Life Technologies), Alpha MEM medium (Thermo), B27 supplements (Thermo), chamber slides (ibidi/ Thermo), Corning® Matrigel® hESC-Qualified Matrix (Sigma), EGF (Thermo), EMEM medium, Fetal calf serum (First Link, PAN Biotec), Heparin (Sigma), FGF2 basic (Thermo), Hyaluronic acid (HA, average molecular weight of 1000KD; Alfa Aesar/Thermo), L-glutamine–penicillin-streptomycin solution (100 X), Sigma), N2 supplements (Thermo), Neurobasal media (Thermo), pyruvate (Life Technologies)phosphate-buffered saline (PBS, Life technologies), Tryple (Thermo), 0.25 wt% trypsin containing 1 mM EDTA (Sigma), ultra-low adhesion U-bottom microplates (Nunc), Amphotericin B 250 µg/ml (Thermo), Glutamine 200mM (Thermo)

#### Cells

The U87 and U251 glioblastoma cell lines were donated by Prof. Kevin Brindle’s laboratory at CRUK, Cambridge. The patient-derived cells, SP20, were given by Dr Richard Mair (Addenbrookes Hospital, Cambridge) as part of the Brain Core program.

#### Glioblastoma cell line U87

*U87 cells were cultured in Alpha MEM complete medium supplemented with 10% FBS, L-Glutamine–Penicillin–Streptomycin solution (1X), in a humidified incubator with 5% CO₂ at 37°C. Upon reaching 70% confluence, cells were detached using trypsin-EDTA and seeded at 5,000 cells in 70 µl per well in 96-well round-bottom ultra-low adhesion microplates. Plates were centrifuged at 1500 rpm for 5 min and incubated with 5% CO₂ at 37°C to facilitate spheroid formation, visible after 48 hours. Spheroids were incubated for 7 days with a media change on day five, then used for migration and proteomic studies*.

#### Glioblastoma cell line U251

U251 cells were cultured in EMEM complete medium supplemented with 15% FBS, L-Glutamine–Penicillin–Streptomycin solution (1X), in a humidified incubator with 5% CO₂ at 37°C. Upon reaching 70% confluence, cells were detached using trypsin-EDTA for 5 minutes at room temperature and seeded at 20,000 cells in 70 µl per well in 96-well round-bottom ultra-low adhesion microplates. Uniformly sized spheroids formed by day 3 and were transferred on day 5 to various concentrations of hyaluronic acid for migration studies.

#### Patient derived primary cells, SP 20

SP20 cells were cultured in serum-free Neurobasal medium supplemented with Glutamine– Penicillin–Streptomycin solution (1X), Amphotericin B (5ml), Glutamine (5 ml) 10 ml B-27, 5 ml N-2, and 20 ng/ml each of EGF, FGF2, and Heparin. T-175 cm² flasks were coated overnight with 10% ECM proteins from Matrigel in PBS, followed by washing with Neurobasal media.

Upon reaching confluence, cells were split using a 1:1 mix of Tryple and Accutase dissociation enzymes and incubated for 5 minutes at room temperature. Cells were seeded at 15,000 in 60 µl per well in 96-well round-bottom ultra-low adhesion microplates. Plates were centrifuged at 1500 rpm for 5 min to assist spheroid formation, visible by day 5. Spheroids were then transferred to various concentrations of hyaluronic acid for migration studies.

#### Preparation of hyaluronic acid matrix for spheroid studies

A stock of 9 wt% HA in MEM alpha complete medium was prepared by thoroughly mixing it to achieve a ‘paste’ texture and left to incubate overnight. Various amounts of this HA stock were added by weight to wells of IBIDI chamber slides and diluted with media to achieve the desired percentages of HA, as in Table 2 below. The chamber slides were incubated overnight in a cell culture incubator to allow them to settle before adding one spheroid per well in 50 µl of media.

**Table 2:** Composition of the HA gels used in this work.

| Final HA wt% | 9 wt% HA paste, mg | Media volume, µl |
| --- | --- | --- |
| 5 | 167 | 83 |
| 3.5 | 117 | 133 |
| 2 | 67 | 183 |

#### Spheroid culture in HA - oxHA mixture

2 wt% HA was prepared as described above. Then 200 mg per well was weighed and allowed to settle inside the well for two hours. Next one spheroid per well in 20 µl of media was transferred and allowed to settle for two hours in a cell culture incubator. Following this 6 wt% oxHA in PBS was prepared, diluted to 2% with Alpha MEM without supplements and 200 µl was added per well.

***Imaging of cells and spheroids*** *was* performed on a Leica DMi1 microscope.

### ^1^H solution-state NMR and T_1_ measurement

All data was acquired using a solid-state Bruker Avance II spectrometer with an operating frequency of 400MHz ^1^H with a double resonance probe (^1^H-^31^P). Hyaluronic acid gels were packed into Kel-F® HR-MAS NMR inserts. Samples containing U87 cells were packed into inserts within 2 hours of fabrication and then stored in the incubator at 37°C, 5% CO_2_ for 24 hours. If needed, samples were centrifuged inside the inserts to remove bubbles. The inserts were then put into a 4mm zirconia rotor and sealed using a Kel-F® rotor cap. All samples were spun at the magic-angle at 5 kHz using compressed air. The spectrometer variable temperature control unit was set to a corrected temperature of 37°C which was calibrated using solid lead nitrate and deuterated methanol with 5 kHz sample spinning at the magic-angle.

T_1_ relaxation was measured using the inversion recovery pulse sequence, with a ^1^H 90° pulse length of 2.5µsec, ^1^H 180° pulse length of 7.140µsec, relaxation delay of 10sec, and the time periods to allow for relaxation between 180° and 90° pulses were: 0.1, 0.2, 0.4, 0.6, 0.8, 1, 1.25, 1.5, 1.75, 2, 2.25, 2.5, 3, 3.25, 3.5, 4, 5, 10, 15, 18, 20sec. Integrated intensities were determined for the signal regions show in Figure 6 below and the resulting signal intensity as a function of relaxation time was fitted according to equation (6) in the SI using MatLab, with rbnmr.

**Figure 6:**
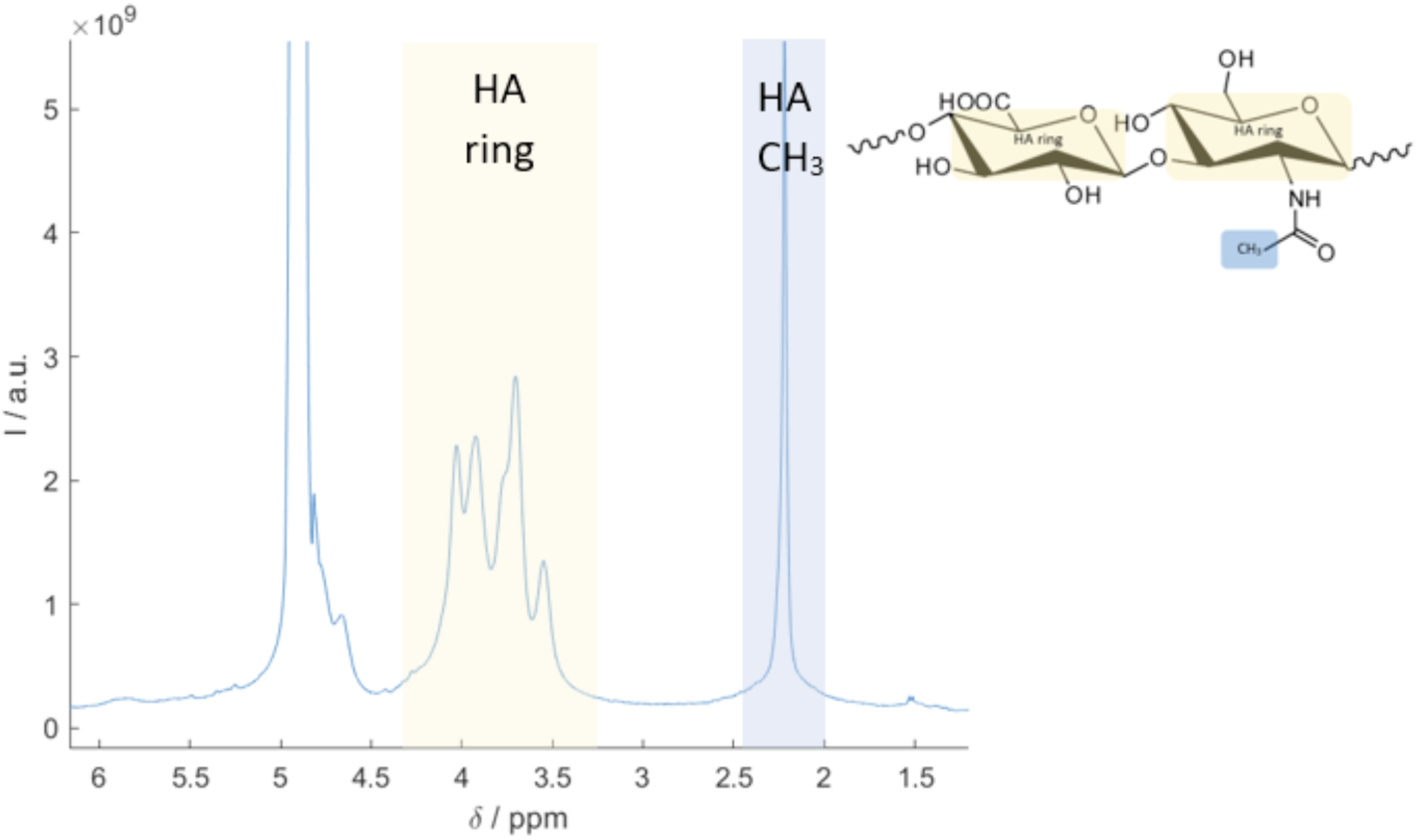
Example ^1^H spectrum of HA gel indicating the signal (integrated) intensities used for T_1_ analysis.

### Rheology

All measurements were carried out using a TA instruments discovery HR2 hybrid rheometer with a 1° 20mm steel conical geometry with the temperature control set to 37°C. Oscillatory time sweeps with a temperature soak time of 60s, experiment time of 60-120s, strain 1% and angular frequency of 10 rad/s were run in between experiments. All experiments were obtained in triplicates unless stated otherwise and the results are presented as an average of three measurements. The following experiments described were run in order. Frequency sweeps were run at constant 1% strain and angular frequency was varied from 0.1 – 100 rad/s. Amplitude sweeps were run at a constant angular frequency of 10 rad/s and varying strain from 0.1 – 500 %. Flow sweeps applied a shear rate of 0.05 – 500 s^−1^ to the samples. For the stress relaxation measurements, if the data was not reproducible after the sample was reloaded following the prerequisite experiments, the samples were reloaded for each individual stress relaxation experiment. The experimental parameters were – temperature soak time of 180s, strain set to 10% for 180s.

### Proteomics

To enable quantitative comparison between proteins expressed in different concentration HMW-HA spheroids were raised simultaneously and planted into the same batch of HA/media/Petri dishes. Protein extraction was performed using the same extraction buffers and according to the same protocol. Protein separation was performed on the same SDS gel, and each band was cut into 5 identical stripes. They were simultaneously digested by trypsin and alkylated. Samples were run on the same LC–MS/MS instrument in the shortest possible time-span between samples to lower down the instrumental variance and instability. Overall, all these precautions ensured that variations in protein detection due to growth conditions, protein extraction, sample handling and instrumentation were minimized. Four protein samples were obtained: (1) spheroid in 5% HMW-HA and (2) total cell dispersion in 2% HMW-HA, both day 7 after spheroid planting; (3) spheroids in 2% HA at the beginning of cell dispersion, day 4 after spheroid planting; (4) spheroid in 2% HA-oxHA mixture, day 4 after spheroid planting. Quantitative comparison was performed for samples 1 and 2, and sample 3 was used as an intermediate point only to corroborate the course of changes. Sample 4, which was obtained on a separate date could be compared to samples 1 and 2 only qualitatively, i.e. by presence or absence of protein of interest.

#### Protein extraction

HA or ox-HA/HA mixture was scooped out of Petri dishes as much as possible without damage to cells/spheroids, which then were transferred into 4-5 fold bigger volume of cOmplete^TM^ Protease Inhibitor Cocktail (Roche, Merck) in PBS and gently mixed for 30 minutes to aid HA dilution. The mixture was split into 50 mL tubes and centrifuged at 4500 rpm at room temperature for 15 min. Supernatant containing HA or oxHA/HA was discarded and another 4-5 volumes of proteinase inhibitor cocktail in PBS was added to the pelleted cells/spheroids still coated in HA. The procedures of mixing and centrifuging were repeated 8 times to completely remove viscose HA smear from pelleted cells/spheroids. Fully cleaned cells/spheroids were combined, snap frozen and ground in liquid nitrogen until fully powdered. The powder was split between low retention Eppendorf tubes and incubated in 10-fold bigger volume of protein extraction buffer (10 wt% TCA, 0.5 wt% DTT (Merck) in pure acetone, 1.5 mL per Eppendorf tube) and left at −20°C overnight. Eppendorf tubes were centrifuged on the next day at 16,000 rpm at 4°C for 20 min. Supernatant was discarded. Pellet was thoroughly mixed with 1 mL of pure acetone per Eppendorf tube and centrifuged at 16,000 rpm at 4°C for 20 min. This procedure was repeated 4 times until pellet became fluffy and soft. After the last centrifugation, acetone was carefully discarded and tubes with pellet were left open in a hood for 3 hours, until pellet was completely dry and acetone smell evaporated. Dried pellet was mixed with Lysis buffer (9M urea, 4 wt% CHAPS, 2% Pharmalyte carrier ampholyte pH 3-10, Merck) and left overnight at room temperature. After centrifugation on the next day at 16,000 rpm at room temperature for 20 min the supernatant containing solubilized proteins was collected, supplemented with up to 1% DTT, snap frozen and kept at −80°C. Protein concentration in the sample was estimated by Pierce^TM^ 660nm protein assay (Thermo) before addition of DTT.

#### LC-MS/MS

15ug of protein in loading buffer was loaded into a Mini-Protean TGX Gels, 4-15%, 10-well, 50ul, bx/10.The gel was run halfway through the gel and stained with Colloidal Coomassie stain. Each sample lane was cut into 6 bands. Bands were transferred into a 96-well PCR plate. The bands were cut into 1mm^2^ pieces, destained, reduced (DTT) and alkylated (iodoacetamide) and subjected to enzymatic digestion with trypsin overnight at 37°C. After digestion, the supernatant was pipetted into a sample vial and loaded onto an autosampler for automated LC-MS/MS analysis.

All LC-MS/MS experiments were performed using a Dionex Ultimate 3000 RSLC nanoUPLC (Thermo Fisher Scientific Inc, Waltham, MA, USA) system and a Q Exactive Orbitrap mass spectrometer (Thermo Fisher Scientific Inc, Waltham, MA, USA). Separation of peptides was performed by reverse-phase chromatography at a flow rate of 300 nL/min and a Thermo Scientific reverse-phase nano Easy-spray column (Thermo Scientific PepMap C18, 2μm particle size, 100A pore size, 75 μm i.d. x 50cm length). Peptides were loaded onto a pre-column (Thermo Scientific PepMap 100 C18, 5μm particle size, 100A pore size, 300 μm i.d. x 5mm length) from the Ultimate 3000 autosampler with 0.1% formic acid for 3 minutes at a flow rate of 15 μL/min. After this period, the column valve was switched to allow elution of peptides from the pre-column onto the analytical column. Solvent A was water + 0.1% formic acid and solvent B was 80% acetonitrile, 20% water + 0.1% formic acid. The linear gradient employed was 2-40% B in 90 minutes. Further wash and equilibration steps gave a total run time of 120 minutes.

The LC eluant was sprayed into the mass spectrometer by means of an Easy-Spray source (Thermo Fisher Scientific Inc.). All *m/z* values of eluting ions were measured in an Orbitrap mass analyzer, set at a resolution of 35000 and was scanned between *m/z* 380-1500. Data dependent scans (Top 20) were employed to automatically isolate and generate fragment ions by higher energy collisional dissociation (HCD, NCE:26%) in the HCD collision cell and measurement of the resulting fragment ions was performed in the orbitrap analyser, set at a resolution of 17500. Singly charged ions and ions with unassigned charge states were excluded from being selected for MS/MS and a dynamic exclusion window of 40 seconds was employed.

#### Database searching

Post-run, the data was initially processed using Protein Discoverer (version 2.5., ThermoFisher). Briefly, all MS/MS data were converted to mgf files and the files were then submitted to the Mascot search algorithm (Matrix Science, London UK, version 2.6.0) and searched against a common contaminants database (125 sequences; 41129 residues); and the UniProt human database (93609 sequences; 37041084 residues). Variable modifications of oxidation (M) and deamidation (NQ) and a fixed modification of carbamidomethyl (C) were applied. Peak areas for each identified peptide were generated and combined to give protein abundance values. The peptide and fragment mass tolerances were set to 20 ppm and 0.1 Da, respectively.

Scaffold (version Scaffold_5.1.1, Proteome Software Inc., Portland, OR) was used to validate MS/MS based peptide and protein identifications. Peptide identifications were accepted if they could be established at greater than 95.0% probability by the Scaffold Local FDR algorithm. Protein identifications were accepted if they could be established at greater than 99.0% probability and contained at least 2 identified peptides. Protein probabilities were assigned by the Protein Prophet algorithm (Nesvizhskii, Al et al Anal. Chem. 2003;75(17):4646-58). Scaffold was also used for comparative quantification: “Quantitative Value (Normalized Toral Precursor Intensity” was selected and normalised to cyclophilin B (PPIB) in each sample to enable quantitative comparison between samples 1, 2 and 3. Protein annotation according to Gene Ontology terms was also performed in Scaffold using NCBI web site: http://www.ncbi.nlm.nih.gov/ and UniProt GOA: http://www.ebi.ac.uk/GOA

