## Supplementary information for "Molecular flexibility of high molecular weight hyaluronic acid measured by NMR has a profound effect on invasion of cancer cells"

### NMR relaxation analysis

For a multispin system, Solomon's equation for the longitudinal relaxation of the macroscopic magnetic moment associated with the *i*th spin, $\bar{I}_{zi}$ can be written as:

$$\frac{d\bar{I}_{zi}}{dt} = -2(\bar{I}_{zi} - \bar{I}_{0i}) \sum_{j \neq i} (W_{1ij} + W_{2ij}) \qquad \text{(i)}$$

$$- (\bar{I}_{zi} - \bar{I}_{zj}) \sum_{j \neq i} (W_{0ij} - W_{2ij}) \qquad \text{(ii)}$$

$$- k_w(\bar{I}_{zi} - \bar{I}_{0i}) + \frac{p_w}{p_{HA}} k_w(\bar{I}_{zw} - \bar{I}_{0w}) \qquad \text{(iii)}$$

Eq 1

where the $W_{aij}$ are the transition probabilities for the *ath*-order quantum transition in a dipolar-coupled $I_{zi}$, $I_{zj}$ spin pair. The first term in equation 1 depends on how far the $\bar{I}_{zi}$ magnetization is from its equilibrium value, $\bar{I}_{0i}$. The second term is driven by differences in the *z*-magnetization between the $I_{zi}$ spin and the surrounding $I_{zj}$ spins to which it is dipolar coupled at any given time. We identify this term as a "spin diffusion" term, because it drives exchange of z-magnetization between the dipolar-coupled *I* spins. This term is zero if all I spins relax at the same rate which will tend to occur if the first term is large, i.e. molecular motion is such that the $\omega_{1ij}$ and $\omega_{2ij}$ transition probabilities are large. The third term comes from exchange of magnetization between HA and water (*w*) $^1$H (through direct $^1$H-$^1$H interaction or chemical exchange), where $p_w$ and $p_{HA}$ are the fraction of $^1$H in the water and HA in the gel respectively and $k_w$ is the effective cross relaxation rate at which water $^1$H magnetization exchanges with HA $^1$H magnetization.

The transition probabilities can be calculated from the equations given in Solomon's original paper [ref] for the case where the *I* spin transitions are driven by dipolar coupling (and molecular motion). Here we consider the case where the molecule undergoes isotropic tumbling with a correlation time $\tau_{macro}$ (= 6 $D_{macro}$ where $D_{macro}$ is the rotational diffusion constant) and internal motion such that the internuclear vector between a pair of $^1$H spins *i* and *j* moves randomly on a cone of cone angle $\theta$ with correlation time $\tau_{int}$. Wallach [ref] has shown that the autocorrelation function *G(t)* in this case is given by:

$$G(t) = \frac{3}{4}\sin^4\theta \exp\left(-\lceil t \rceil \left(\frac{1}{\tau_{macro}} + \frac{4}{\tau_{int}}\right)\right) +$$
$$3\sin^2\theta\cos^2\theta \exp\left(-\lceil t \rceil \left(\frac{1}{\tau_{macro}} + \frac{1}{\tau_{int}}\right)\right) +$$
$$\left(\frac{1}{2}(3\cos^2\theta - 1)\right)^2 \exp\left(-\lceil t \rceil \left(\frac{1}{\tau_{macro}}\right)\right)$$

Eq 2

Following Solomon[1] Harris,[2] and Navon and Lanir,[3] this results in the transition probabilities:

$$W_{0ij} = (2\pi R)^2 \frac{1}{20}\left[\frac{3}{2}\,\tau_{c1}\sin^4\theta + 6\,\tau_{c2}\sin^2\theta\cos^2\theta + \frac{1}{2}\tau_{macro}\,(3\cos^2\theta - 1)^2\right]$$

$$W_{1ij} = (2\pi R)^2 \frac{3}{40}\left[\frac{3}{4}\sin^4\theta\left(\frac{2\tau_{c1}}{1+\omega_0{}^2\tau_{c1}{}^2}\right) + 3\sin^2\theta\cos^2\theta\left(\frac{2\tau_{c2}}{1+\omega_0{}^2\tau_{c2}{}^2}\right) + \left(\frac{1}{2}(3\cos^2\theta - 1)\right)^2\left(\frac{2\tau_{macro}}{1+\omega_0{}^2\tau_{macro}{}^2}\right)\right]$$

$$W_{2ij} = (2\pi R)^2 \frac{3}{10}\left[\frac{3}{4}\sin^4\theta\left(\frac{2\tau_{c1}}{1+4\omega_0{}^2\tau_{c1}{}^2}\right) + 3\sin^2\theta\cos^2\theta\left(\frac{2\tau_{c2}}{1+4\omega_0{}^2\tau_{c2}{}^2}\right) + \left(\frac{1}{2}(3\cos^2\theta - 1)\right)^2\left(\frac{2\tau_{macro}}{1+4\omega_0{}^2\tau_{macro}{}^2}\right)\right]$$

Eq 3

where

$$R = \left(\frac{\mu_0}{4\pi}\right)\frac{\gamma_i\gamma_j}{r_{ij}}\left(\frac{\hbar}{2\pi}\right)$$

$$\frac{1}{\tau_{c1}} = \frac{1}{\tau_{macro}} + \frac{4}{\tau_{int}}$$

$$\frac{1}{\tau_{c2}} = \frac{1}{\tau_{macro}} + \frac{1}{\tau_{int}}$$

In our plots in Figure 2, we have plotted $2(W_{1ij} + W_{2ij})$ (the "molecular motion" term) and $(W_{0ij} - W_{2ij})$ (spin diffusion term) as a function of θ and $\tau_{int}$ for a fixed $\tau_{macro}$, the amplitudes of the two components in eq 1. Additional plots are in Fig S1.

The relaxation times for a given $^1H$ spin *i* is governed by the sum of all its pairwise interactions with neighbouring $^1H$ spins (eq 1), which will typically have different $\theta$ and *R* for each pair for a given internal motion in the HA molecule that involves the *i* spin. On top of this, there will be many different internal molecular motional modes, likely with different $\tau_{int}$ and different distribution of θ for the $^1H$-$^1H$ internuclear vectors. Our aim here is to find the values of θ and $\tau_{int}$ for which the effect of the internal molecular motion on the $^1H$ spin-lattice relaxation time can be expected to dominate over the spin diffusion term, and so to describe the necessary internal motions that must be present for HA molecules where spin diffusion is demonstrably quenched. (Note that the molecular motion term in Eq 1 is multiplied by $(\bar{I}_{zi} - \bar{I}_{0i})$, the difference between the current *i* spin *z*-magnetization and its equilibrium value whilst the spin diffusion term is multiplied by $(\bar{I}_{zi} - \bar{I}_{zj})$, the difference between the *i* and *j* spin *z*-magnetizations. When *i* and *j* relax with similar rates, this latter term is small, and thus the spin diffusion term is multiplied by a relatively small cofactor. Where the *i* spin *z*-magnetization is far from equilibrium, the molecular motion term is multiplied by a relatively large factor.)

The coupling between HA and water $^1$H spin-lattice relaxation introduced by the third term of eq 1 means that plots of $\bar{I}_{zi}(t)$ v *t,* are fitted as a sum of two exponential curves with apparent relaxation rate constants $T_{1i}^{+}$ and $T_{1i}^{-}$:[4]

$$\bar{I}_{zi}(t) = c_i^{+} \exp(-t/T_{1i}^{+}) + c_i^{-} \exp(-t/T_{1i}^{-})$$

Eq 4

where

$$c_i^{+} + c_i^{-} = 1$$

and the true *i* spin $T_{1i}$ given by:

$$\frac{1}{T_{1i}} = \frac{c_i^{+}}{T_{1i}^{+}} + \frac{c_i^{-}}{T_{1i}^{-}}$$

Eq 5

In the inversion recovery experiment used in this work, the signal intensity $S_i(t)$ from the *i* $^1$H at time *t* is given by:

$$S_i(t) = S_0\big(1 - 2\bar{I}_{zi}(t)\big)$$
$$= S_0\big(1 - 2(c_i^{+} \exp(-t/T_{1i}^{+}) + c_i^{-} \exp(-t/T_{1i}^{-}))\big)$$

Eq 6

The values given below and in Figure 2 for the HA methyl and ring $^1$H $T_1$ are calculated from eq 5 from the values of $c_i^{\pm}$ and $T_{1i}^{\pm}$ (*i* = methyl or ring) determined by fitting each experimental spin-lattice curve $S_i(t)$ v *t* as a sum of two exponentials according to eq 6, the fits giving the values in Table S1 and the $S_i(t)$ v *t* plots in Fig S2.

**Determination of the degree of oxidation in oxHA**

The integrated intensity of signals in the $^1$H NMR of oxHA is proportional to the number of protons contributing to each respective signal. In the case of HA, for the NH-CO-$CH_3$ group methyl, the protons (three per methyl group) in this signal will have contributions from to both oxidised and non-oxidised HA, as oxidation should leave these signals almost unchanged, so the integrated intensity of this peak represents both species. After oxidation the 2' and 3' groups on the glucuronic acid subunit of HA form aldehydes, both of which appear as the broad signal at 9.2 ppm; in aqueous solution, the aldehydes will be in equilibrium with their hydrated forms which give rise to the three peaks in the 4.9 - 5.25 ppm region. Thus, the integrated intensity of these three signals plus that of the signal at 9.2 ppm are proportional to the aldehyde concentration in the sample. Using the equation and integrated intensity values in the figure below to calculate the molar ratio between signals arising from the two hydrated and non-hydrated aldehyde protons versus signal intensity arising from the HA + oxHA methyl $^{1}$Hs, the degree of oxidation can be calculated as roughly 35%.

OxHA was further characterized by FTIR, revealing the expected aldehyde signal at 1730 $cm^{-1}$:[5]

The time scale of oxHA crosslinking with cell culture media amine groups was determined by solution-state NMR as described in figure S5.

### Supplementary Figures

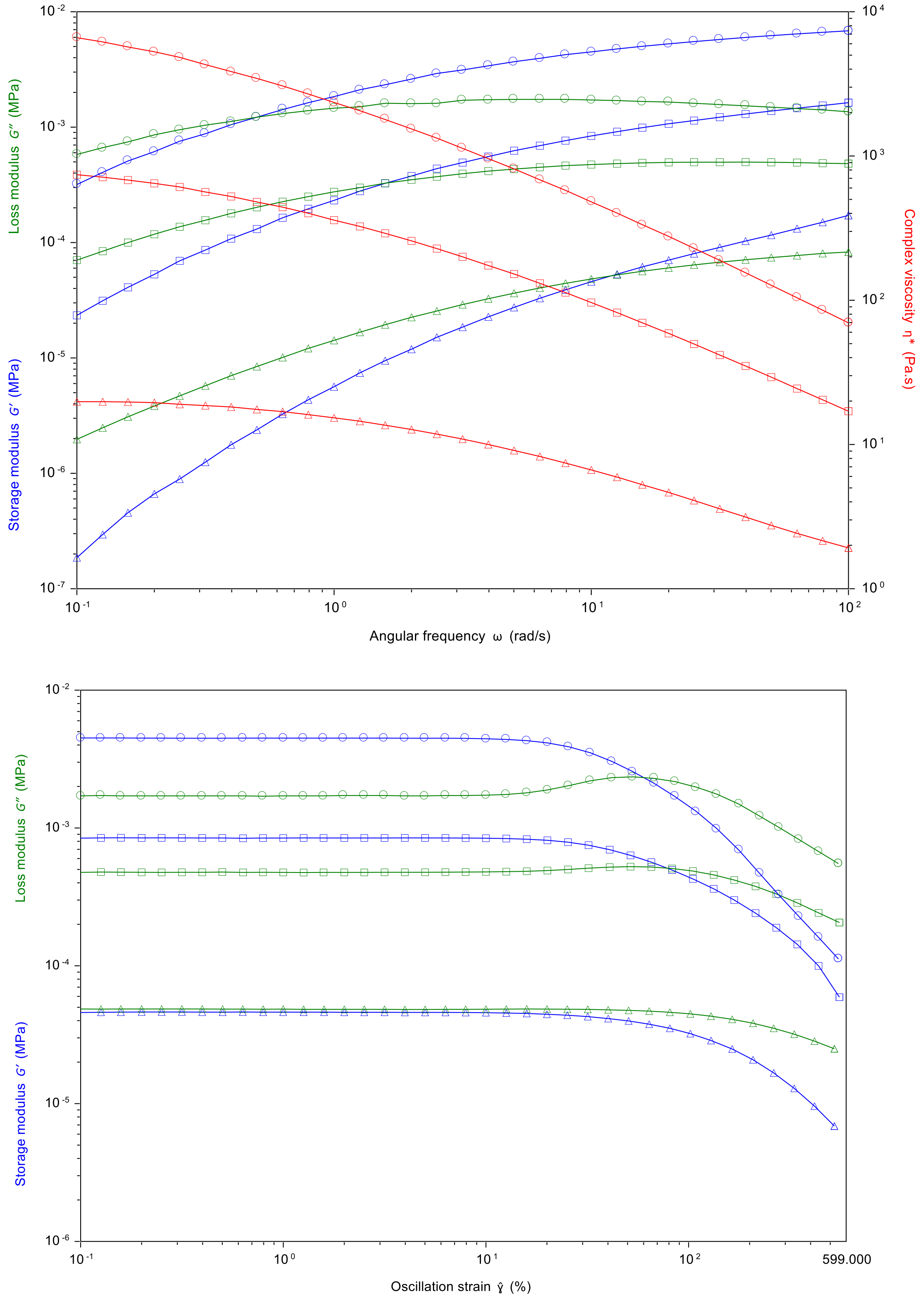


**Figure S1:** Rheology data for the HMW-HA gels used in this work (circles, 10 wt% HA; squares, 5 wt% HA; triangles, 2 wt% data).

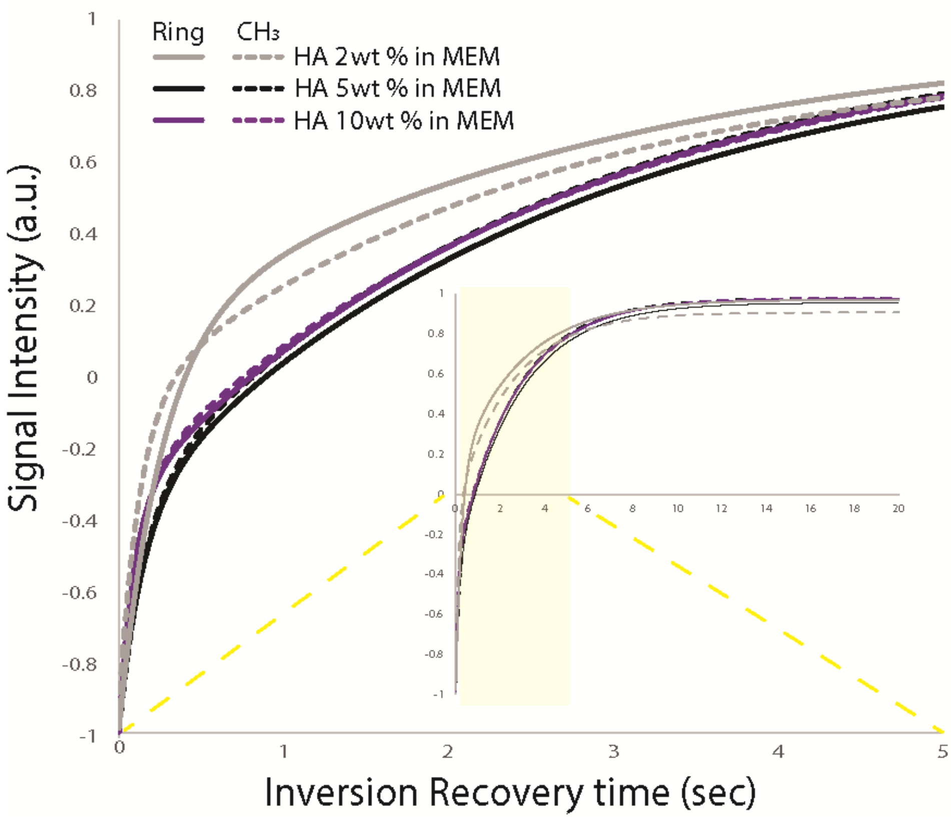


**Fig S2:** Plots of the fitted inversion recovery $^1$H signal intensity for different concentration of HA for the HA methyl and ring non-OH $^1$H signals, fitted according to equation (6) above. The ring and methyl $^1$H relaxation curves are highly similar to each other in 5 and 10 wt% HA indicating that spin diffusion dominates the $^1$H spin-lattice relaxation mechanism at these HA concentrations. In contrast, the methyl and ring $^1$H relaxation curves for 2 wt% HA are significantly different, indicating that spin diffusion is not the dominant mechanism at this lower HA concentration.

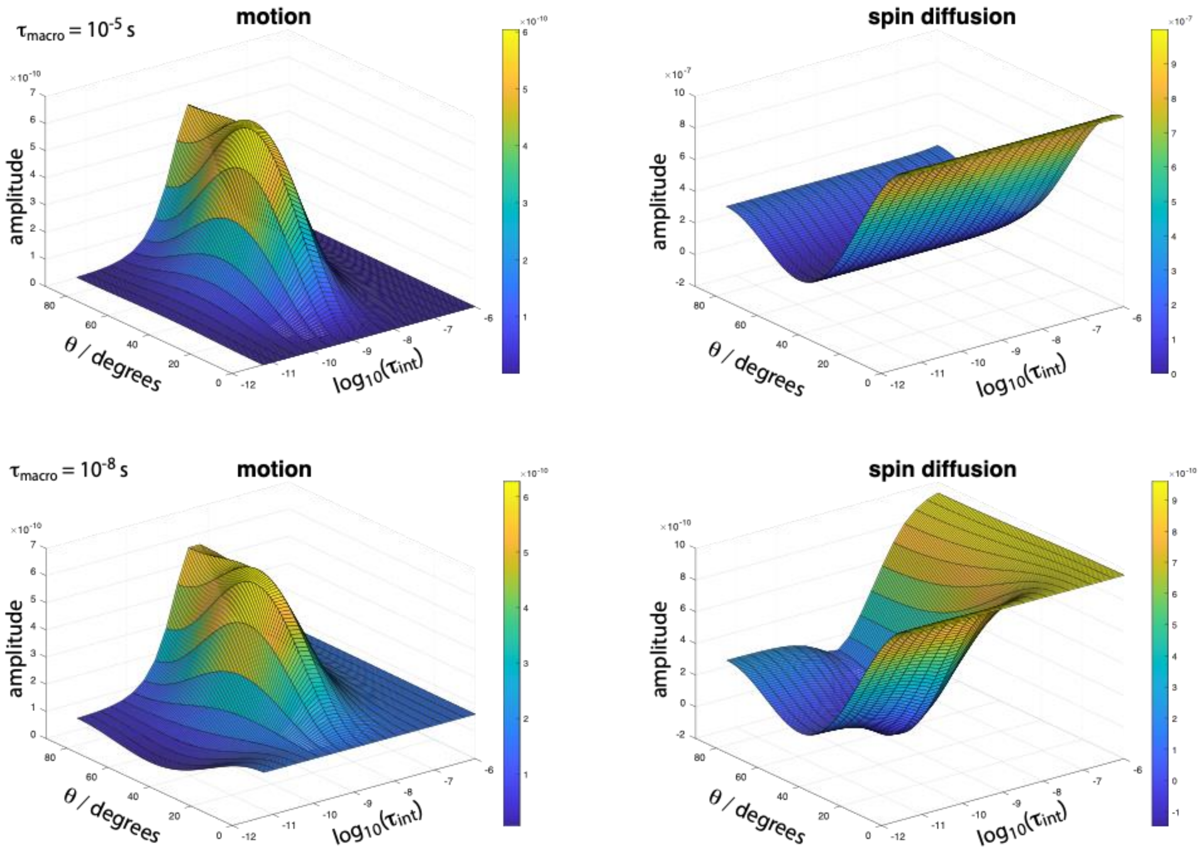


**Figure S3:** Surface plots showing how the amplitudes of terms (i) (motion) and (ii) (spin diffusion) in equation 1 above vary with the correlation time for internal motion on a cone of cone angle $\theta$ and correlation time $\tau_{int}$, for isotropic tumbling correlation times $\tau_{macro}$ as given on the plots.

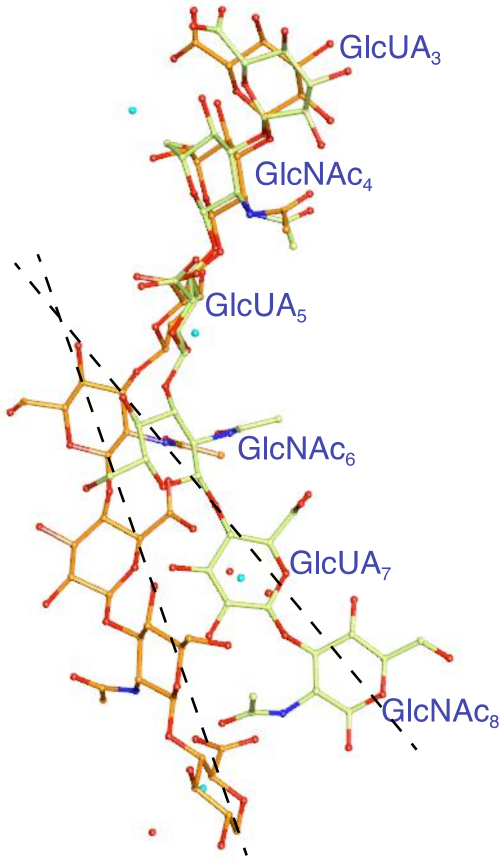


**Figure S4:** Taken from reference[6]. The conformation of HA (oligomers) bound to CD44 (green) is distorted from that for free HA (orange, structure taken from 3HYA). Dotted lines show the approximate distortion of the HA backbones between the structures.

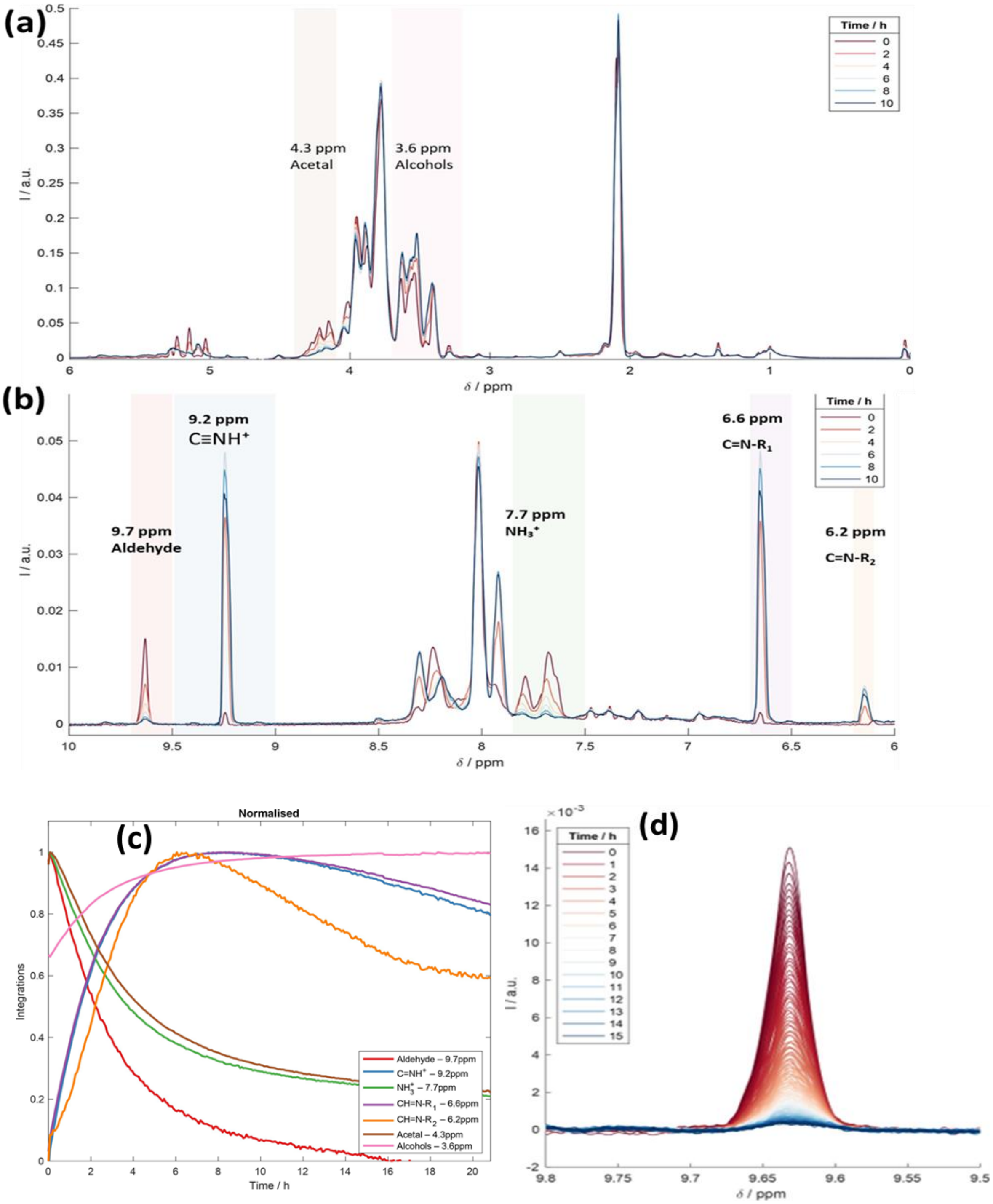


**Figure S5:** $^{1}$H solution-state NMR spectra of 2 wt% oxHA in Alpha MEM media. (a) and (b) show $^{1}$H signals corresponding to aldehydes at 9.7 ppm, CH=N groups at 9.2 ppm, acetals around 4.3 ppm, and alcohols around 3.6 ppm. (c) shows the intensity of the indicated signals as a function of time after mixing oxHA and media. The decay of signal intensity observed in the characteristic aldehyde peak at 9.7 ppm over time, as illustrated in (d), indicates that the aldehyde groups undergo crosslinking reactions with $NH_2$ groups of amino acids and serum proteins in media. Other peaks at 7.7 ppm and 4.3 ppm, which correspond to $NH_3^+$ and acetal groups respectively, also exhibit intensity decay. These data confirms the covalent crosslinking of oxHA with proteins and / or amino acids in media. Similar crosslinking is expected in the extracellular matrix proteins deposited by the cells in the culture.

### Supplementary tables

#### Table S1: Fitted $^1H$ $T_1$ relaxation time constants

| **HA wt%** | **$^1H$** | **% component** | **$T_1$ / s** |
|---|---|---|---|
| **2 wt%** | $CH_3$ | 53.4 | 2.43 |
| | | 46.6 | 0.13 |
| | ring | 44.4 | 2.85 |
| | | 55.6 | 0.25 |
| | water | 55.8 | 4.23 |
| | | 44.2 | 0.07 |
| **5 wt%** | $CH_3$ | 68.1 | 2.56 |
| | | 31.9 | 0.15 |
| | ring | 69.5 | 2.64 |
| | | 30.5 | 0.15 |
| | water | 78.5 | 2.89 |
| | | 21.5 | 0.03 |
| **10 wt%** | $CH_3$ | 66.0 | 2.57 |
| | | 34.0 | 0.10 |
| | ring | 67.9 | 2.57 |
| | | 32.1 | 0.09 |
| | water | 71.0 | 2.88 |
| | | 29.0 | 0.03 |
